# Functional and transcriptomic analyses in *Neurospora crassa* reveal the crucial role of N-glycoprotein deglycosylation process in fungal homeostasis

**DOI:** 10.64898/2026.03.25.714127

**Authors:** Anastasios Samaras, Tanim Jabid Hossain, Magnus Karlsson, Georgios Tzelepis

## Abstract

N-glycosylation is an essential post-translational modification required for proper protein folding, stability, trafficking, and secretion in eukaryotes. In such organisms, an efficient endoplasmic reticulum (ER) quality control, such as the ER-associated degradation (ERAD) pathway, is critical for maintaining cellular homeostasis. During ERAD, terminally misfolded glycoproteins undergo N-deglycosylation prior to proteasomal degradation, a process typically mediated by peptide N-glycanase (PNGase). However, in the filamentous fungi, the PNGase seems to be catalytically inactive, indicating evolutionary divergence from the canonical PNGase pathway. Filamentous fungi also encode endo-β-N-acetylglucosaminidases (ENGases), particularly members of glycoside hydrolase family 18 (GH18), which may compensate for the loss of canonical PNGase activity. Here, we investigated the roles of the cytosolic GH18 ENGase and a putative acidic PNGase in *N. crassa* using transcriptomic and functional approaches. Our results demonstrate that the cytosolic GH18 ENGase is an active deglycosylating enzyme likely associated with the ERAD pathway, whereas no deglycosylation activity was detected for the acidic PNGase. Deletion of the ENGase severely compromises tolerance to diverse stress conditions and induces substantial transcriptomic reprogramming, including upregulation of a GH20 exo-β-N-acetylhexosaminidase under ER stress. These findings identify cytosolic ENGase as a key component of fungal proteostasis and suggest that *N. crassa* activates alternative compensatory mechanisms to maintain protein quality control when canonical deglycosylation pathways are impaired.

## Introduction

N-glycosylation is one of the most crucial post-translational modifications in eukaryotic proteins, as it is essential for proper protein folding, stability, solubility, intracellular trafficking, and efficient secretion of extracellular enzymes [1]. Filamentous fungi secrete large quantities of N-glycoproteins, including hydrolytic enzymes, cell wall– remodeling factors, and stress-responsive proteins, making the integrity of the secretory pathway critical for fungal physiology [2, 3].

N-linked glycosylation is initiated in the endoplasmic reticulum (ER) through the co-translational transfer of a Glc₃Man₉GlcNAc₂ oligosaccharide to asparagine residues within the conserved Asn–X–Ser/Thr motif by the oligosaccharyltransferase complex [1]. Following transfer, the glycan undergoes sequential trimming by ER glucosidases I and II, generating monoglucosylated intermediates that serve as quality-control signals for entry into the calnexin/calreticulin folding cycle [4, 5]. These early glycan-processing events allow misfolded glycoproteins to be recognized, reglucosylated by UDP-glucose:glycoprotein glucosyltransferase, and repeatedly engaged with lectin chaperones until a native conformation is achieved [6]. However, when N-glycoproteins repeatedly fail to fold correctly, they are exported to the cytosol and degraded by the proteasome via the endoplasmic reticulum–associated degradation (ERAD) pathway [7, 8]. During this process, removal of N-glycans by deglycosylating enzymes is essential for further process, as proteasomes are unable to efficiently degrade all glycosylated polypeptides [9].

In the N-deglycosylation process two enzymes play a crucial role: the peptide N-glycanase (PNGase) and the endo-β-N-acetylglucosaminidases (ENGases) [10]. PNGase encoded by *png1* in yeasts and animals, catalyzes the removal of N-glycans from glycoproteins by hydrolyzing the amide bond between the innermost N-acetylglucosamine (GlcNAc) and asparagine residues, thereby preparing misfolded substrates for proteasomal degradation [11]. In *Saccharomyces cerevisiae*, Png1 physically associates with Rad23 to facilitate the delivery of deglycosylated polypeptides to the 26S proteasome, directly linking deglycosylation to ER-associated degradation (ERAD) efficiency [12].

In contrast, the PNG1 ortholog in *Neurospora crassa* is catalytically inactive due to conserved substitutions within the essential Cys–His–Asp catalytic triad, indicating a functional divergence from the canonical PNGase role [13]. Despite lacking enzymatic activity, *N. crassa* PNG1 is essential for hyphal polarity, since deletion of *png-1* results in severe polarity defects and cell lysis, suggesting a structural or scaffolding role independent of deglycosylation [13]. This evolutionary shift, loss of catalytic PNGase activity, while retaining essential cellular functions, appears to be conserved among many filamentous ascomycetes, implying strong selective pressure to maintain the non-enzymatic functions of PNG1 [14]. Furthermore, PNG1 in *N. crassa* is genetically uncoupled from rad23, unlike in yeast where Png1 and Rad23 form a functional complex, providing additional evidence for divergence in the fungal PNGase system [13]. Notably, a second gene encoding a putative acidic PNGase has been identified in some filamentous species genomes, however, its precise biological role remains unclear [13, 14, 15].

Fungal ENGases, belonging to glycoside hydrolase (GH) family 18, represent a unique class of enzymes in filamentous fungi capable of removing or processing N-glycans and may partially compensate for the loss of canonical PNGase activity [16]. Filamentous fungal genomes often encode two fungi-specific GH18 ENGases, including one predicted cytosolic and one secreted enzyme [14]. In addition, a second putative cytosolic ENGase belonging to GH85, exhibiting both transglycosylating and hydrolytic activity, has been identified in some filamentous fungal species [14, 17, 18].

Previous studies have demonstrated that cytosolic GH18 ENGases play critical roles in multiple aspects of fungal biology, including secretion, stress tolerance, and antagonistic interactions [19, 20]. However, the broader contribution of N-deglycosylation pathways to fungal physiology remains poorly understood. In this study, we investigate the roles of the cytosolic GH18 ENGase and the acidic PNGase in *Neurospora crassa* using transcriptomic and functional approaches. Our data show that the cytosolic GH18 ENGase is an active deglycosylating enzyme possibly involved in the ERAD pathway, in contrast to the acidic PNGase, for which no such activity was observed. Furthermore, deletion of this ENGase severely compromises fungal tolerance to diverse stress conditions and induces significant transcriptomic changes, including the upregulation of a GH20 exo-β-N-acetylhexosaminidase under ER stress. These findings suggest that the cytosolic ENGase plays a crucial role in fungal homeostasis and that *N. crassa* can activate alternative pathways to compensate for disruptions in protein quality control machineries.

## Results and Discussion

### The gh18-10 ENGase is an active cytosolic de-glycosylating enzyme

As it is known that the cytosolic PNGase (PNG-1) in *N. crassa* is enzymatically inactive [13], we focused our studies on three other putative de-glycosylating enzymes: the putative cytosolic and secreted ENGases gh18-10 and gh18-11 respectively and the acidic PNGase (pngA), previously identified in the *N. crassa* genome [13, 14]. To investigate the in vitro enzymatic activity of these enzymes, we heterologously expressed them in *png1Δ S. cerevisiae* cells since this species genome lacks other de-glycosylating enzymes [21]. Western blot analysis targeting the V5 epitope tag confirmed the successful expression of gh18-10, gh18-11, and pngA in yeast cells (Fig. S1A). Furthermore, we investigated whether the proteins were N-glycosylated by treating them with the endoglycosidase (Endo)-H enzyme. Our results showed a shift in the molecular weight of gh18-11 and pngA after Endo-H treatment (Fig. S1B), indicating that these two proteins are highly N-glycosylated and possibly secreted as it was previous predicted. In contrast, no band shift was observed for gh18-10, suggesting that it is not N-glycosylated, further confirmed its predicted cytosolic localization. These results are consistent with our previous data showing that the putative secreted ENGase in *Trichoderma atroviride* undergoes extensive N-glycosylation, unlike the cytosolic one. This suggests that N-glycosylation is crucial for the folding and functionality of this enzyme in adverse extracellular environments. [13].

Since these proteins have been confirmed to be heterologously expressed in yeast cells, the next step was to investigate the in vitro de-glycosylation activity of these enzymes. To that end, the highly N-glycosylated enzyme S-alkylated RNase B was incubated with yeast cell extracts expressing the three studied putative de-glycosylating enzymes. Our results showed that the cytosolic gh18-10 ENGase is an active de-glycosylating enzyme, since a clear shift in the substrate’s molecular weight was observed (Fig. 1A). In contrast, no shift in mobility was observed in S-alkylated RNase B treated with yeast cell extracts expressing the gh18-11 ENGase and the acidic PNGase pngA, indicating that these enzymes are not active de-glycosylating enzymes under these studied conditions (Fig. 1A). Together with our previous studies in *T. atroviride* [13], these results further support the role of the cytosolic GH18 ENGase as a substitute for the enzymatically inactive PNGase in the de-N-glycosylation machinery of filamentous ascomycetes.

**Fig. 1.**
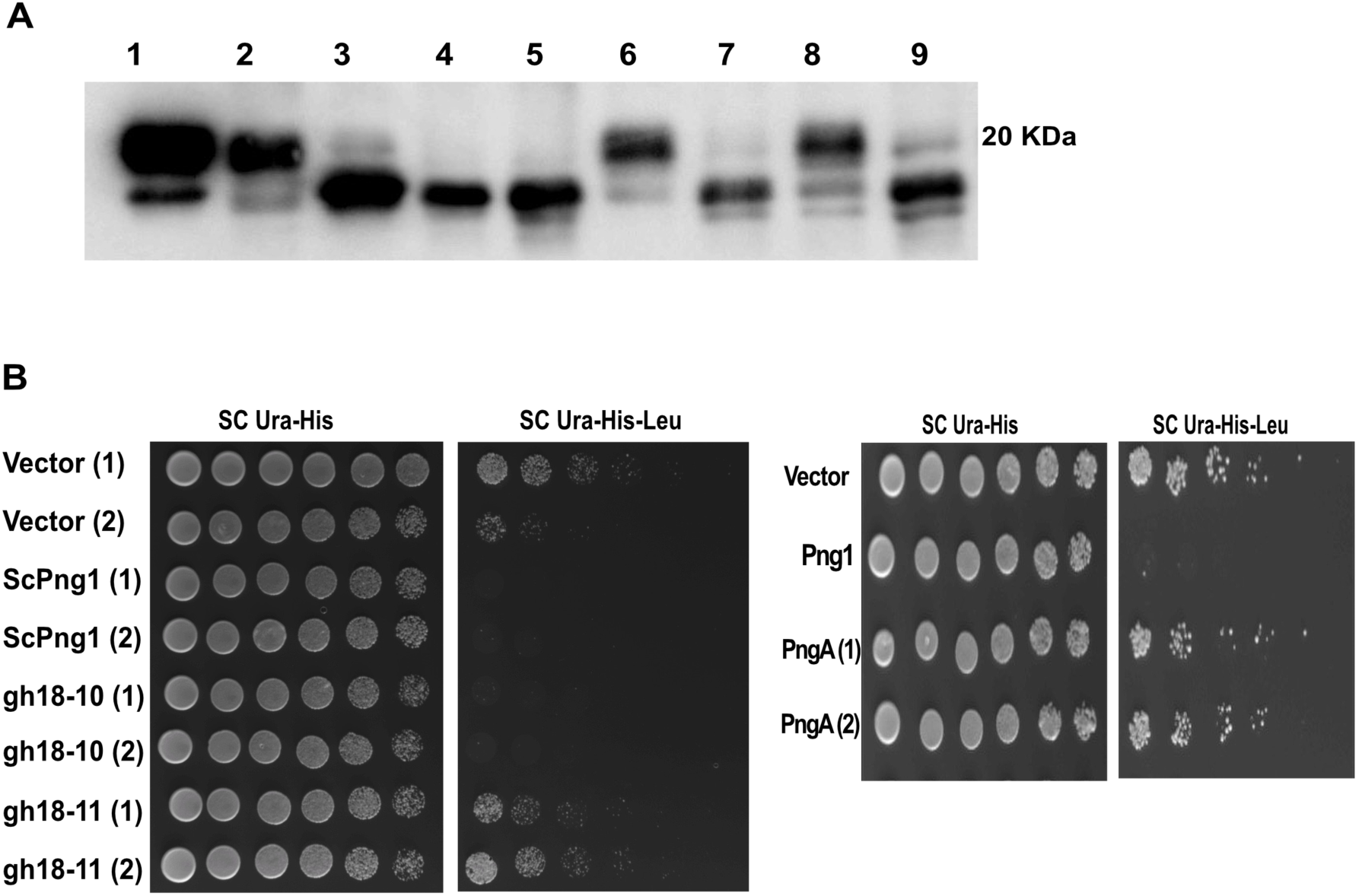
The cytosolic gh18-10 ENGase is an active deglycosylated enzyme possibly involved in the ERAD process. (**A**) In vitro deglycosylation activity of *Neurospora crassa* ENGases and PNGase enzymes. *Neurospora crassa* gh18-10, gh18-11 and pngA were expressed in *png1*Δ (A) *S. cerevisiae* cells, and proteins were extracted from each strain and incubated with S-alkylated RNase B. Deglycosylation activity resulted in decreased molecular weight and was detected by a shift in migration on SDS–PAGE gel, by immunoblotting using anti-RNase B antibody and HRP-conjugated secondary antibody. (**1)** S-alkylated RNase B + lysis buffer, (**2)** S-alkylated RNase B + empty vector, (**3)** S-alkylated RNase B + PNGase F, (**4)** S-alkylated RNase B + gh18-10, **(5)** S-alkylated RNase B + Eng18A + PNGase F, (**6**) S-alkylated RNase B + gh18-11, (**7**) S-alkylated RNase B + gh18-11 + PNGase F, (**8**) S-alkylated RNase B + pngA, (**9**) S-alkylated RNase B + pngA + PNGase F. (**B**) Effect of gh18-10, gh18-11 and pngA on RTL stability. Enzymes were co-expressed with the RTL protein in *S. cerevisiae png1*Δ cells. Six-fold serial dilutions of each cell suspension were spotted onto SC-Ura-His + 2% galactose plate and to SC-Ura-His-Leu + 2% galactose plate. Yeast strains expressed only the empty vector (pRS316) were used as a negative control exhibiting a defective ERAD, while the Png1 complemented yeast strains (ScPng1) were used as a control exhibiting efficient degradation of RTL.

### The gh18-10 ENGase is implicated in the ERAD pathway

We next examined whether these enzymes can functionally replace cytoplasmic PNGase in the ERAD pathway of *S. cerevisiae*. To address this question, we employed the in vivo RTL assay as previously described [22]. RTL is a chimeric membrane protein composed of a luminal RTA domain, the transmembrane domain of Pdr5, and a cytoplasmic Leu2 domain [22]. The RTL assay allows the detection of ERAD defects based on yeast growth. In strains carrying a leu2 mutant allele, growth in leucine-deficient medium depends on the accumulation of Leu2 supplied by an intact RTL fusion protein; thus, growth in the absence of leucine occurs only when RTL degradation is impaired [22].

Our data show that yeast cells heterologously expressing the gh18-10 ENGase failed to grow on minimal medium, indicating that this enzyme promotes RTL degradation in a manner similar to *S. cerevisiae* Png1 (Fig. 1B). These results suggest a role for this cytosolic ENGase in the ERAD process, consistent with previous observations for homologous ENGases in *T. atroviride* [16]. In contrast, yeast cells heterologously expressing the gh18-11 ENGase grew on leucine-deficient medium at levels comparable to cells carrying the empty vector (Fig. 1B), indicating that these enzymes are unable to degrade this ERAD substrate. Additionally, we investigated whether the acid PNGase could degrade the RTL protein. Acidic PNGases have sporadically been identified in filamentous ascomycete genomes, but they have no significant sequence similarities with cytosolic PNGases [14, 23]. Our data showed that yeast cells that heterologously expressed the pngA enzyme were unable to degrade the RTL protein, similar to gh18-11 (Fig. 1B). This indicates that this enzyme cannot replace the inactive cytosolic PNGase in *N. crassa*. These data, combined with the widespread presence of cytosolic GH18 ENGases in ascomycete genomes, further support the conserved enzymatic role of this enzyme in filamentous fungal N-de-glycosylation machinery.

### Deletion of *gh18-10* alters fungal stress tolerance

Our previous studies have demonstrated that cytosolic ENGases play a crucial role in filamentous fungal morphology and physiology, affecting growth, conidiation, response to abiotic stress conditions, and protein secretion [19, 20]. In this study, we further expanded the response of *N. crassa* gh18-10 and pngA deletion strains to variable stress conditions, including oxidative, anoxic, endoplasmic reticulum (ER), and cell wall stresses. Consequently, the radial growth of *gh18-10* and *pngA* mutants was compared to the wild type (WT). No statistical differences were observed among the three strains under control or under cell wall (caffeine) and DTT stress conditions (p > 0.05, Fig. 2A, B). Interestingly, DTT disrupts disulfide bond formation in the ER, but no phenotype was observed. This implies that cytosolic de-N-glycosylation may be more relevant to glycoprotein quality control after ER export, rather than directly modulating the ER folding machinery under reductive stress.

**Fig. 2.**
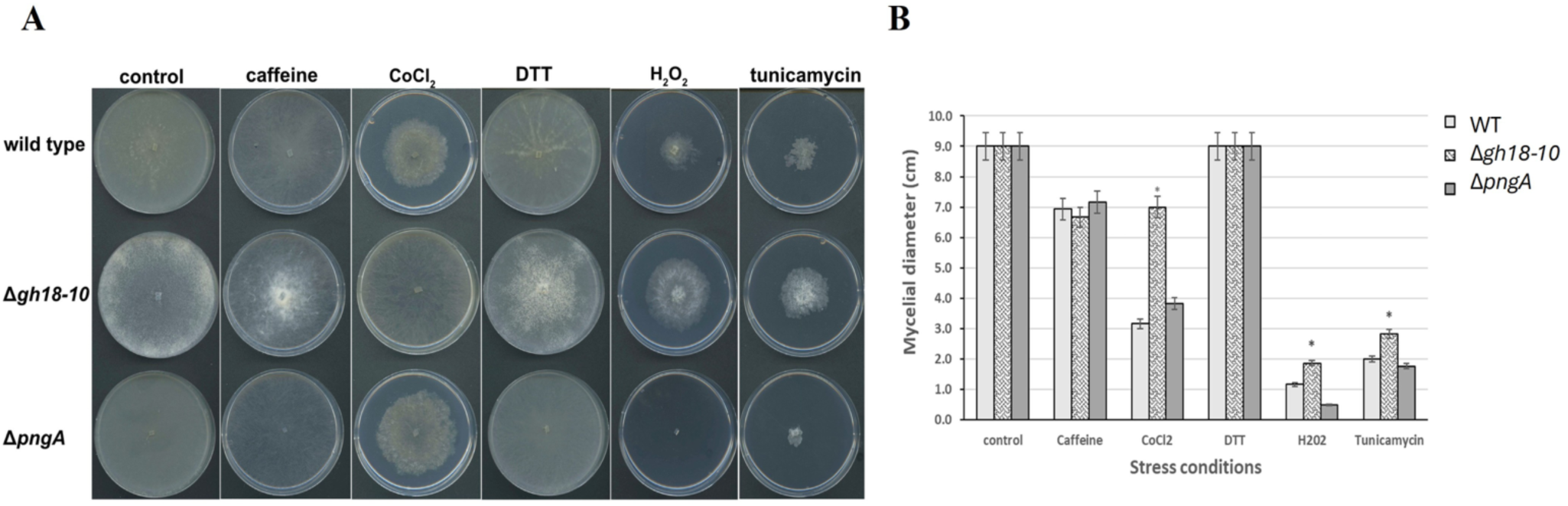
Deletion of the *gh18-10* ENGase increases *Neurospora crassa* tolerance to variable stress conditions. (**A**) Representative colony morphology of *Neurospora crassa* wild type (WT), Δ*gh18-10*, and Δ*pngA* strain grown on solid VM medium under control conditions or supplemented with caffeine (cell wall stress), CoCl₂ (hypoxia), DTT (ER stress), H₂O₂ (oxidative stress), or tunicamycin (ER stress). (**B**) Mycelia growth was quantified as mycelial diameter (cm) under each condition (right) 24 hours post inoculation at 30°C. Bars represent the mean ± error. Asterisks indicate statistically significant differences relative to WT (*p* < 0.05) according to Tukey test.

However, significant statistical differences were observed under specific stress treatments. The *gh18-10* deletion strain exhibited significantly increased mycelial growth compared to the wild type (WT) under CoCl₂-induced anoxia, H₂O₂-mediated oxidative stress, and tunicamycin-induced endoplasmic reticulum (ER) stress (p < 0.05; Fig. 2A, B). In contrast, the *pngA* deletion strain displayed reduced tolerance relative to both the WT and the *gh18-10* mutant under H₂O₂ and tunicamycin treatments (p < 0.05; Fig. 2A, B). Tunicamycin is well known to induce ER stress by inhibiting N-glycosylation, resulting in the accumulation of misfolded proteins within the ER lumen. Additionally, tunicamycin activates the cell wall integrity (CWI) signaling pathway, highlighting the functional cross-talk between ER proteostasis mechanisms and cell wall maintenance systems [24, 25]. This interaction is particularly critical in filamentous fungi, where polarized hyphal growth relies on tightly coordinated protein secretion and continuous cell wall remodeling [26, 27]. The enhanced growth of the *gh18-10* mutant under ER, oxidative, and anoxic stress conditions suggests that loss of cytosolic ENGase activity may alter the degradation dynamics of misfolded glycoproteins. Such alterations could promote adaptive stress signaling or reduce the accumulation of toxic degradation intermediates. Furthermore, disruption of de-N-glycosylation may induce a proteostatic imbalance that primes stress-response pathways, including oxidative stress and unfolded protein response (UPR) signaling, thereby conferring a growth advantage under adverse conditions. Further, the increased tolerance to both anoxic and oxidative stress also indicates a broader stress adaptation extending beyond ER-specific perturbations, potentially reflecting altered redox homeostasis or improved management of oxidative damage. Overall, these findings suggest that gh18-10 functions as a negative regulator of stress tolerance under specific abiotic stress conditions.

### Deletion of *gh18-10* has a negative impact in *N. crassa* sexual reproduction

Since previous studies reported a reduced number of protoperithecia in the *gh18-10* deletion strain [19], we further examined whether this deletion affects sexual reproduction. To address this, crosses between different *N. crassa* strains were performed and the production of viable ascospores was assessed. Our results demonstrated that deletion of *gh18-10* causes pronounced defects in sexual development (Fig. 3A,B). In crosses involving the Δ*gh18-10* strain, perithecia were either absent or, when formed, were largely empty and incapable of producing ascospores (Fig. 3A). This phenotype was especially evident when Δ*gh18-10* served as the protoperithecial parent, as ascospore production was consistently undetectable. Quantification of ascospore output from multiple pairwise crosses with other mutant strains supported this trend, whereas other strain combinations, including those involving the Δ*pngA* strain, yielded detectable ascospore numbers (Fig. 3B). Collectively, these findings further support our previous observations that disruption of the major de-N-glycosylation activity in *N. crassa* severely impairs sexual reproduction. Given the importance of glycoprotein processing for cell wall remodeling, signaling, and differentiation, loss of gh18-10 likely disrupts the maturation or function of key N-glycosylated proteins required for ascogenous tissue formation and ascospore development. The comparatively mild phenotype of Δ*pngA* further supports the notion that gh18-10 represents the major de-N-glycosylation activity necessary to sustain sexual reproduction.

**Fig 3.**
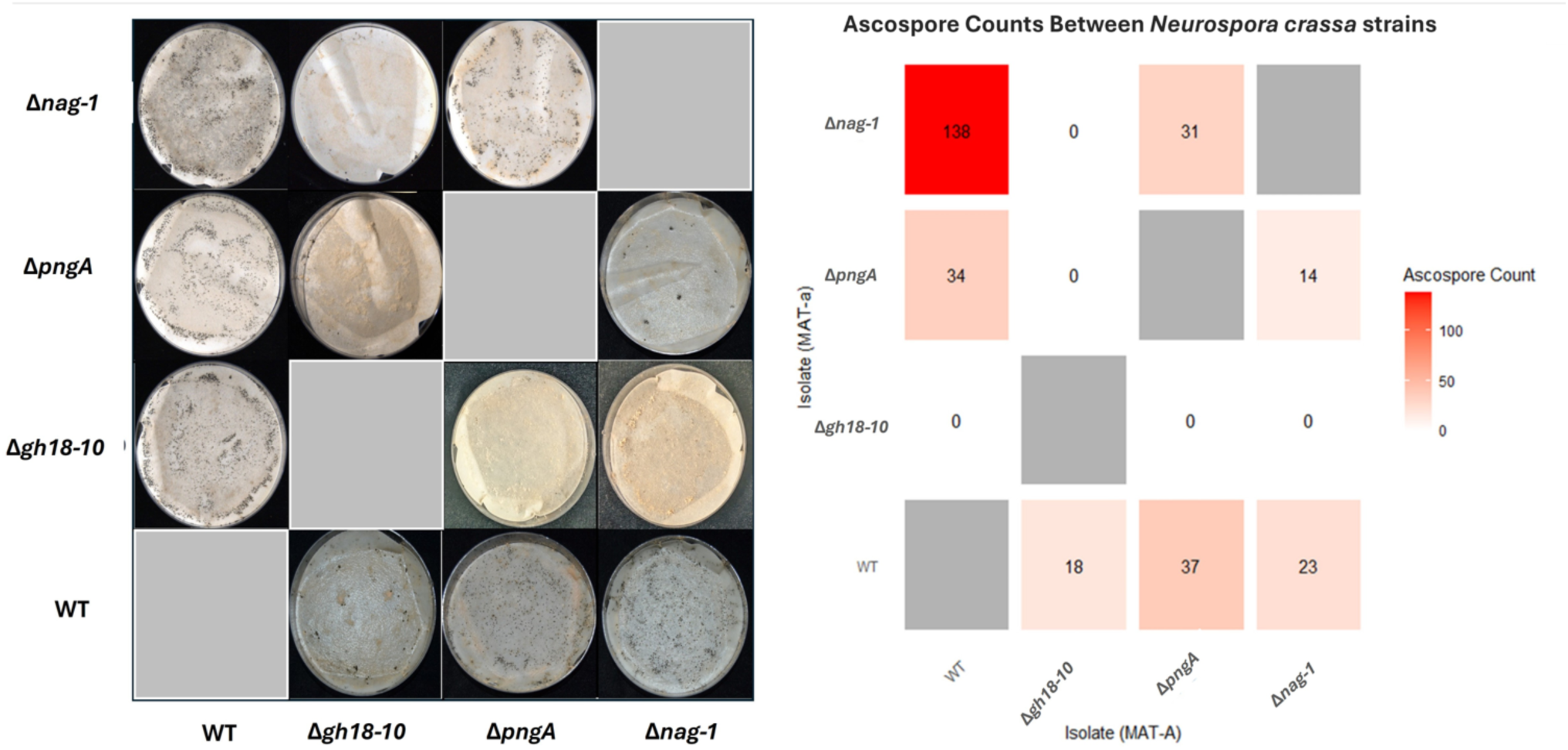
Deletion of the *gh18-10* ENGase impairs *Neurospora crassa* sexual reproduction. (**A**) representative images of perithecial development and ascospore formation from reciprocal crosses between wild type (WT), Δ*nag-1*, Δ*pngA*, and Δ*gh18-10* strains of opposite mating types grown under standard crossing conditions 14 days after crossing. Rows correspond to MAT-a isolates and columns to MAT-A isolates. (**B**) heatmap summarizing ascospore counts obtained from the corresponding crosses. Numbers within each cell indicate the total number of ascospores per ml recovered per cross, with color intensity reflecting ascospore abundance.

### Deletion of *gh18-10* leads to significant transcriptome alterations

In order to investigate the potential role of de-N-glycosylation in fungal physiology we performed a comprehensive transcriptomic analysis of *N. crassa* strains. As an initial step, we assessed whether deletion of de-N-glycosylation–related enzymes caused major transcriptomic alterations under non-stress conditions. To this end, the transcriptomes of the *gh18-10* and *pngA* deletion strains were compared with that of the wild type (WT) during growth on Vogel’s medium in the absence of stress agents. Our analysis revealed that the transcriptome of the *gh18-10* deletion strain differed markedly from that of WT under control conditions (Fig. 4A). In total, 1,471 genes were differentially expressed (−2 < log₂ fold change < 2), with 553 genes upregulated and 918 genes downregulated relative to WT (Fig. 4B). This extensive transcriptional remodeling indicates that loss of *gh18-10* strongly affects global gene expression even in the absence of external stress. In contrast, the *pngA* deletion strain exhibited a much more limited transcriptional response under the same conditions, with only 76 differentially expressed genes, including 45 upregulated and 31 downregulated genes (Fig. 4B). This pronounced difference in the number of affected genes is consistent with the PCA results and further supports the conclusion that *gh18-10* deletion has a substantially greater impact on the basal transcriptome than deletion of *pngA*. Heatmap visualization further highlights these differences, revealing a broad and coordinated shift in gene expression in the *gh18-10* mutant compared to WT, whereas the *pngA* mutant displays only minor changes (Fig. S2A). Notably, differentially expressed genes in the *gh18-10* strain cluster into distinct expression modules, suggesting coordinated regulation of multiple functional pathways (Fig. S2B). These data indicate that gh18-10 likely has a central homeostatic function in *N. crassa*.

**Fig. 4.**
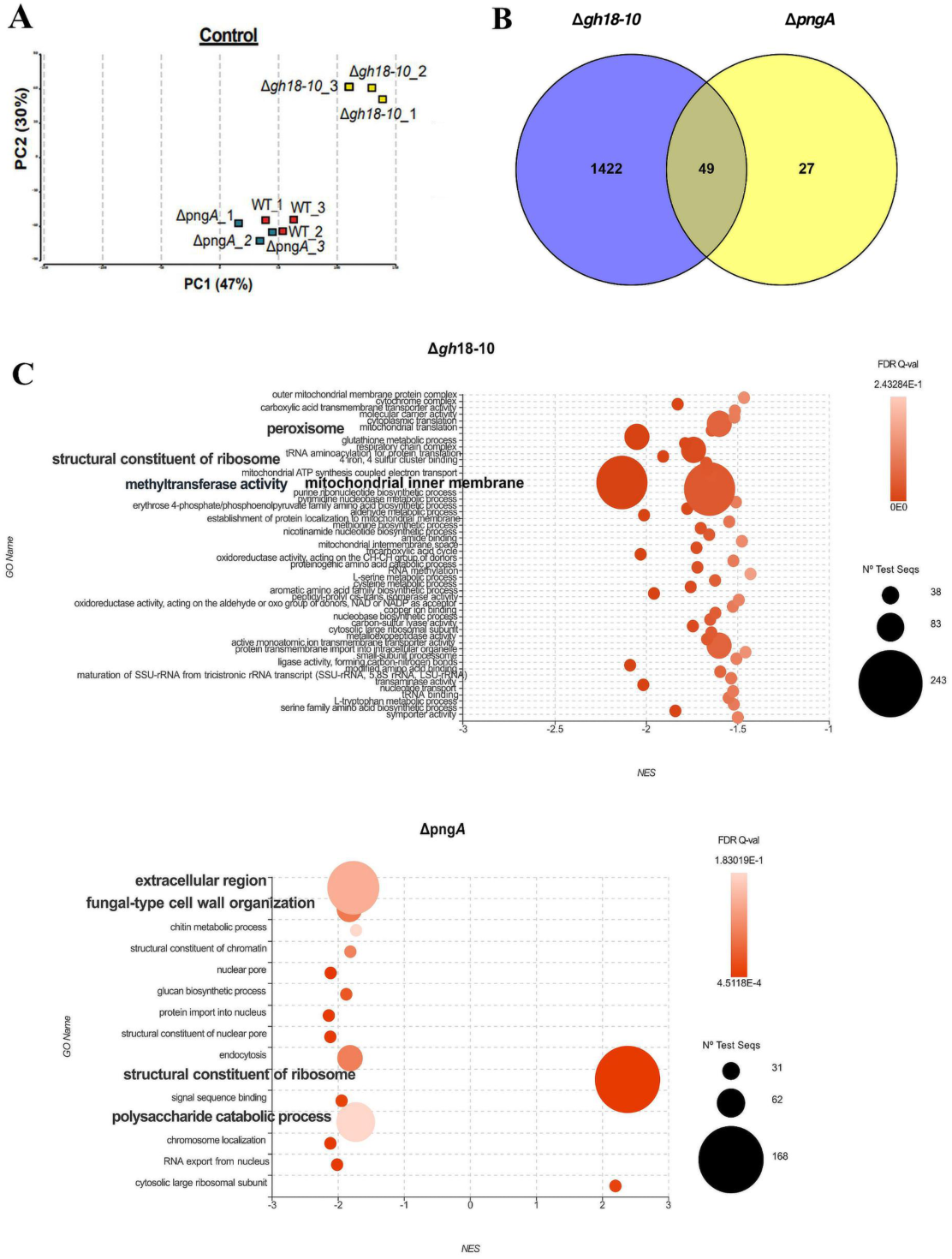
Deletion of the *gh18-10* ENGase causes broad alternations in *Neurospora crassa* transcriptome. (A) PCA plot of normalized transcriptome profiles from *N. crassa* WT, Δ*pngA*, and Δ*gh18-10* strains grown in liquid VM media (control conditions). (B) Venn diagram of differentially expressed genes (DEGs) of Δ*pngA*, and Δ*gh18-10* grown under control conditions as normalized to the WT transcriptome. (C) Bubble plot of Gene Ontology (GO) enrichment for DEGs under control conditions on Δ*gh18-10* and Δ*pngA* strains. Bubble size indicates the number of genes associated with each GO term.

Gene Ontology (GO) enrichment analysis under the same control conditions revealed that the two mutant strains affect distinct functional categories relative to WT. In the Δ*gh18-10* strain, the most strongly enriched GO terms were predominantly associated with intracellular metabolism and organelle-related functions, with a pronounced enrichment of mitochondrial categories, including mitochondrial inner membrane, cytochrome/respiratory chain complexes, and mitochondrial translation (Fig. 4C). Additional enriched terms included peroxisome-related functions, methyltransferase activity, and a wide range of metabolic and biosynthetic processes, such as glutathione metabolism, TCA cycle and other carboxylic acid–related processes, amino acid biosynthesis and metabolism, as well as RNA methylation and translation-associated terms, including structural constituents of the ribosome (Fig. 4C). In contrast, the Δ*pngA* strain showed enrichment of functional categories linked primarily to cellular interfaces and trafficking processes. These included the extracellular region and fungal-type cell wall organization, with prominent representation of chitin metabolic and glucan biosynthetic processes, along with polysaccharide catabolism. Enriched terms also encompassed several nucleus- and transport-related categories, such as nuclear pore organization, protein import into the nucleus, RNA export from the nucleus, and endocytosis (Fig. 4C).

In addition, we specifically examined GO terms associated with sexual reproduction and completion of the sexual cycle. This analysis indicated that the sexual developmental transcriptional program is already attenuated under control conditions. Affected genes included those involved in fruiting body development and maturation (e.g., *NCU02925*, annotated as “fruiting body”), female sexual development (e.g., *NCU09915*, *female sexual development-1*), and meiosis-related processes (e.g., *NCU01510* and *NCU08064*, annotated with meiotic GO terms) (Fig. S3). Overall, these findings imply that gh18-10 is a key regulator of basal cellular physiology in *N. crassa*, influencing mitochondrial activity, metabolism, and development, whereas pngA appears to contribute more specifically to cell wall dynamics and transport processes.

### Deletion of *gh18-10* causes broad transcriptome remodeling during stress

As previously demonstrated, deletion of *gh18-10* results in pronounced alterations in *N. crassa* phenotype to various abiotic stresses. To elucidate the potential mechanisms by which de-N-glycosylation contributes to this phenotype, a comprehensive transcriptomic analysis was performed on fungal deletion strains subjected to ER, oxidative, and anoxic stress conditions. These stress agents were selected because they elicited the most pronounced phenotypic differences among the strains in our assays. Principal component analysis (PCA) was conducted to evaluate global similarities and differences in transcriptomic profiles across genotypes and treatment conditions. In all PCA plots, biological replicates clustered tightly, demonstrating the high reproducibility and robustness of the RNA-seq data (Fig. 5A). Notably, the *gh18-10* deletion strain displayed a transcriptomic profile that was clearly distinct from those of both the wild-type (WT) and *pngA* mutant strains under all tested conditions. In contrast, transcriptomic profiles of the WT and **pngA** deletion strains clustered relatively closely together (Fig. 5A).

**Fig. 5.**
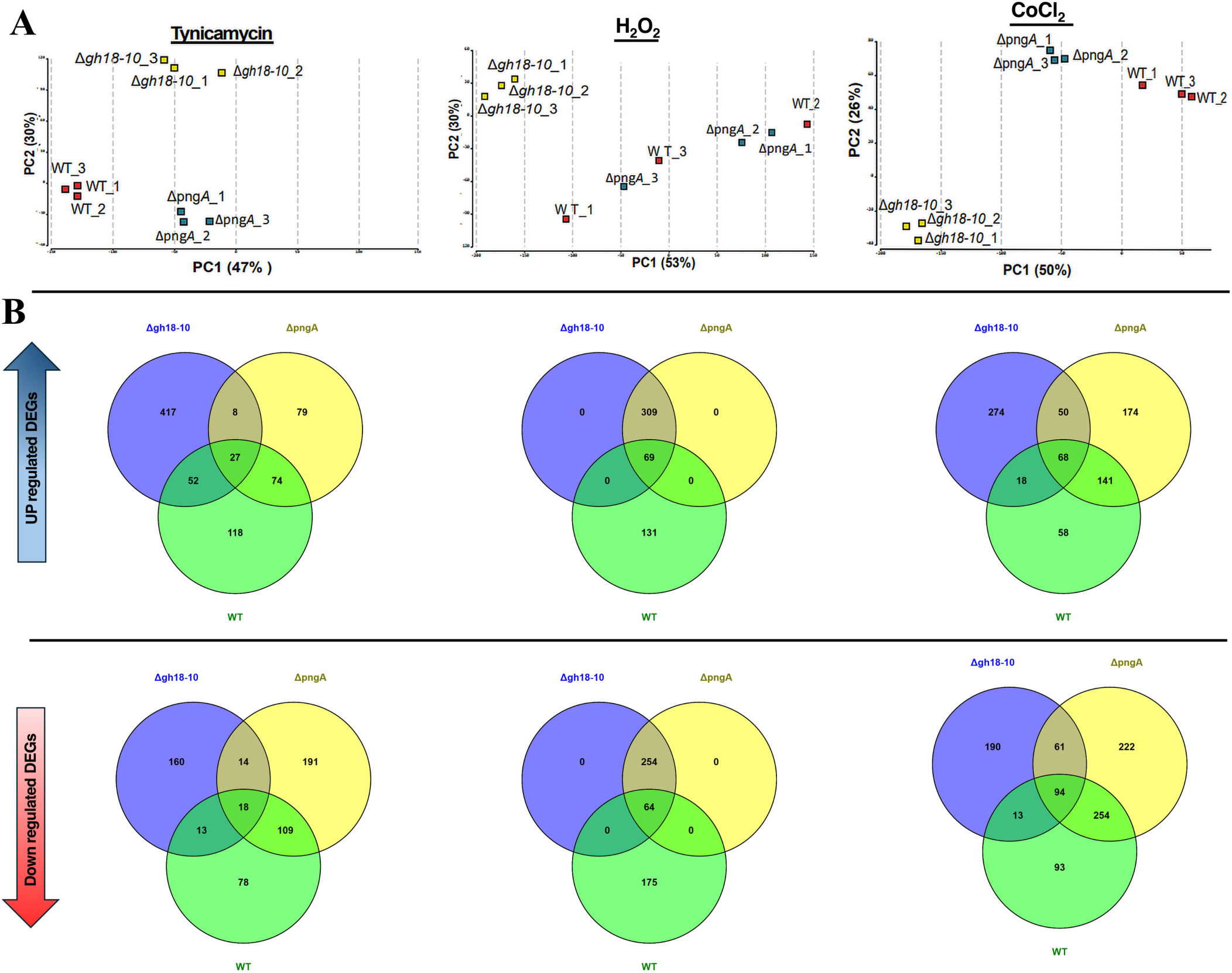
Deletion of *gh18-10* alters *Neurospora crassa* transcriptome under stress conditions. (A) PCA plots of normalized transcriptome profiles of *N. crassa* WT, Δ*pngA*, and Δ*gh18-10* under variable abiotic stress conditions. (B) Venn diagrams of uniquely up and down regulated genes under variable stress condition. *N. crassa* Δ*gh18-10* and Δ*pngA* and WT strains grown in liquid Vogels media treated with tunicamycin (ER-stress), H2O2 (oxidative stress) and CoCl2 (hypoxic stress).

Across the three stress conditions, differential gene expression (DEG) patterns revealed both shared and genotype-specific transcriptional responses (Fig. 5B). Under tunicamycin treatment, Δ*gh18-10* exhibited the largest mutant-specific induction, with 417 uniquely upregulated genes. In contrast, Δ*pngA* showed only 79 uniquely induced genes, while the WT had 118; just 27 genes were commonly induced across all three strains (Fig. 5B). Tunicamycin-mediated repression was more pronounced in Δ*pngA* (191 uniquely downregulated genes) than in Δ*gh18-10* (160), with 18 genes consistently downregulated in all strains (Fig. 5B). Under oxidative stress (H₂O₂), neither mutant displayed strain-specific DEGs in either direction. Instead, the two mutants shared large DEG sets that were distinct from WT (309 commonly upregulated and 254 commonly downregulated genes), whereas WT maintained a substantial number of unique responses (131 upregulated and 175 downregulated) (Fig. 5B). A smaller subset of genes was shared among all three strains (69 upregulated and 64 downregulated) (Fig. 5B). In the presence of CoCl₂, both mutants again demonstrated pronounced strain-specific transcriptional changes. For upregulated genes, Δ*gh18-10* had 274 unique genes, Δ*pngA* had 174, and WT had 58 (Fig. 5B). For downregulated genes, Δ*gh18-10* showed 190, Δ*pngA* had 222, and WT had 93. In addition, there were notable shared responses: 68 genes were upregulated and 94 downregulated across all three strains. The two mutants also exhibited considerable overlap, with 50 genes co-upregulated and 61 genes co-downregulated (Fig. 5B).

Given that endoplasmic reticulum (ER) stress plays a central role in the accumulation of misfolded glycoproteins in eukaryotic cells, we further investigated fungal responses to impaired de-N-glycosylation by focusing on the *N. crassa* transcriptome following exposure to tunicamycin (TNC). Under TNC-induced stress, 402 genes were uniquely upregulated in the Δ*gh18-10* strain compared with the WT and ΔpngA strains, as well as relative to the *Δgh18-10* control transcriptome (Fig S4A). The overlap between the Δ*gh18-10* TNC transcriptome and its untreated control was limited to only 15 genes, indicating that the majority of induced genes represent a TNC-specific response rather than baseline transcriptional differences (Fig. 6A). A similar pattern was observed for downregulated genes. The Δ*gh18-10*_TNC transcriptome contained 159 uniquely downregulated genes, with minimal overlap with WT_TNC (13 genes) or Δ*pngA*_TNC (14 genes). A comparable number of differentially expressed genes (15 genes) was shared among Δ*gh18-10*_TNC, WT_TNC, and Δ*pngA*_TNC. Notably, only one gene overlapped exclusively between Δ*gh18-10*_TNC and the Δ*gh18-10* control transcriptome on the downregulated side (Fig. S4A).

**Fig. 6.**
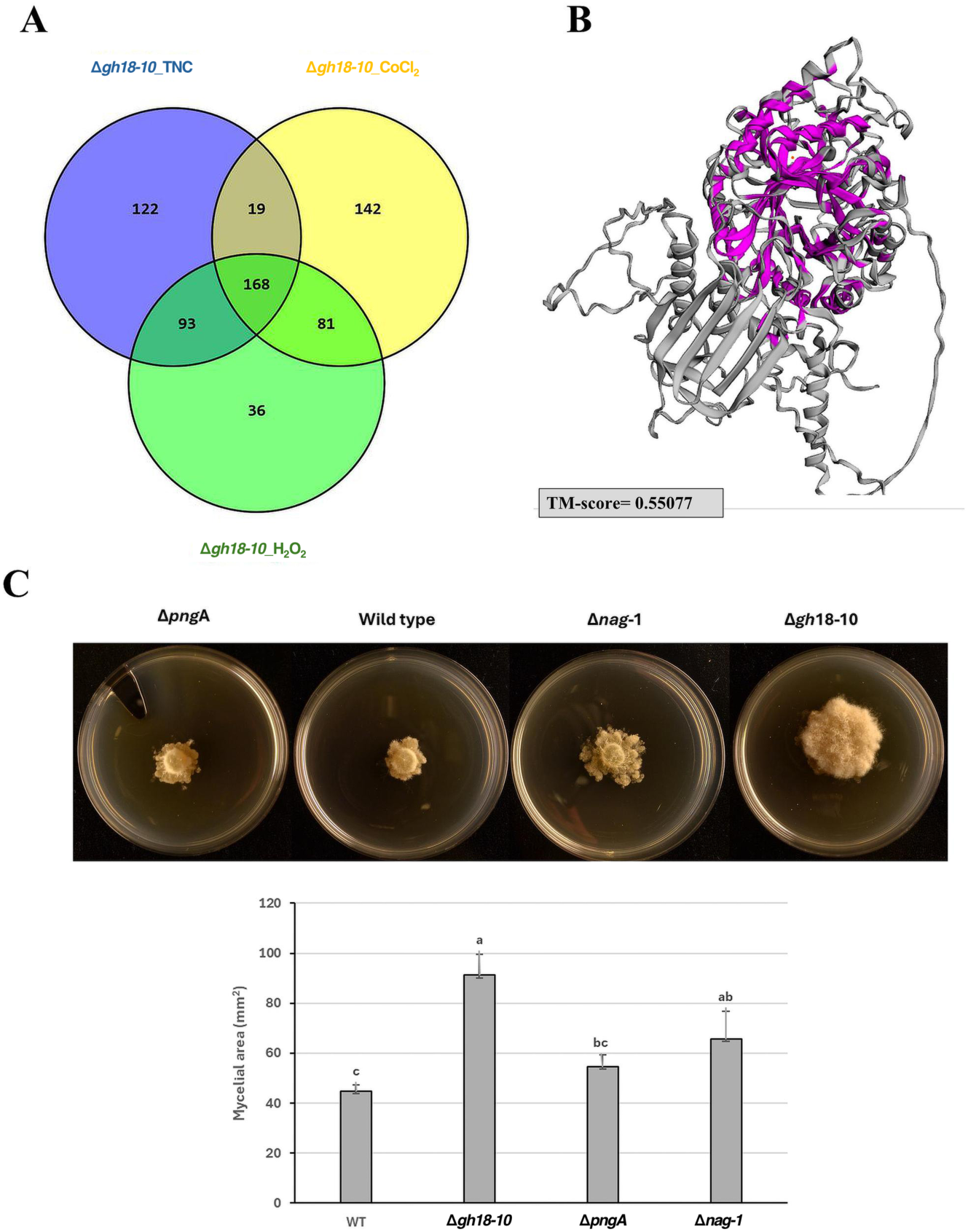
The *Neurospora crassa nag-1* gene coding for a putative GH20 N-acetyl hexosaminidase is uniquely induced on *gh18-10* deletion strain upon ER stress. **(A**) Venn diagram of differentially expressed genes (DEGs) of Δ*pngA*, Δ*gh18-10* and WT strain grown under tunicamycin ER stress conditions. (**B**) Predicted schematic overview and structural comparison highlighting similarities between the nag-1 and gh18-10 enzymes. PDB structures were generated using AlphaFold 3, and structural alignments were performed with TM-align using default parameters. (**C**) Phenotypic analysis of *nag-1* deletion strain under tunicamycin stress conditions. Mycelial radial growth was measured 24 h of incubation at 30 °C. Different letters (a, b, c) indicate statistically significant differences, and bars represent the mean ± error. Asterisks indicate statistically significant differences (*p* < 0.05) according to Tukey test.

Weighted gene co-expression network analysis (WGCNA) further supported the DEG-based findings, demonstrating that the TNC response is concentrated within a limited number of co-expression modules and varies by genotype. The highest DEG enrichment was observed in the blue and turquoise modules, with Δ*gh18-10* exhibiting the strongest overall signal. Module eigengene analysis clearly separated TNC-treated samples from controls. Modules such as blue, turquoise, brown, and pink showed increased eigengene expression under TNC treatment, whereas green and yellow modules exhibited the opposite trend. These patterns indicate coordinated, module-level transcriptional programs that are differentially activated across genotypes (Fig. S4B). A comprehensive list of all differentially expressed genes (DEGs) across all treatments and strains, including gene identifiers, fold changes, and statistical significance values for each contrast, is provided in Supplementary File 1. Overall, these data suggest that gh18-10–mediated de-N-glycosylation plays a critical regulatory role in maintaining ER homeostasis and proteostasis under stress. Its deletion leads to exaggerated and highly specific transcriptional reprogramming, particularly under ER stress, likely reflecting impaired processing or clearance of misfolded glycoproteins and consequent activation of compensatory stress-response pathways.

### ER stress induces GH20 N-acetylhexosaminidase gene expression in the Δ*gh18-10* strain

We next focused on genes uniquely upregulated in Δ*gh18-10* during TNC exposure to determine whether alternative glycan-processing activities are induced in the absence of the *gh18-10* gene. Comparison of upregulated DEG sets in Δ*gh18-10* across TNC, H₂O₂, and CoCl₂ treatments revealed a substantial shared core of 168 genes, together with pronounced condition-specific subsets. Specifically, 122 genes were uniquely induced by TNC, 142 by CoCl₂, and 36 by H₂O₂ (Fig. 6A). Functional enrichment analysis of the TNC-specific upregulated genes was performed using g:Profiler. The most significantly enriched Gene Ontology (GO) terms were predominantly associated with transporter and trafficking functions, including bicarbonate/oxalate transmembrane transporter activity, bicarbonate/oxalate transport, and plus-end-directed vesicle transport along microtubules (Fig. S5A). Notably, no clearly annotated deglycosylation-related terms were detected at the GO level. This absence prompted a manual inspection of the induced gene set. Manual curation identified several genes encoding enzymes with plausible roles in glycan processing. Among these was *NCU10852* (*nag-1*), annotated as a β-hexosaminidase (N-acetylhexosaminidase; GH20), along with additional glycan-related proteins, including a putative glycosyltransferase and other carbohydrate-active enzymes (Fig. S5B).

Given that NAGases hydrolyze terminal N-acetylglucosamine residues from glycoconjugates [28], the induction of *nag-1* represents a biologically plausible candidate for partial functional compensation of glycan-processing capacity under TNC treatment in the Δ*gh18-10* background. Supporting this hypothesis, protein structural alignment revealed measurable similarity between nag-1 and gh18-10 (Fig. 6B). Phenotypic characterization further indicated functional relevance. Colony growth assays under TNC treatment demonstrated that the Δ*nag-1* strain exhibited greater radial mycelial growth compared with WT, with a phenotype more closely resembling that of Δ*gh18-10* (Fig. 6C). Quantification of mycelial area confirmed that Δ*nag-1* formed significantly larger colonies than WT, while no statistical differences were observed between Δ*nag-1* and Δ*gh18-10*, (p > 0.05) indicating enhanced vegetative growth in the absence of the NAGase gene, resembling the effect seen when *gh18-10* is deleted. (Fig. 6D). These data indicate that both genes likely participate in related glycan-processing pathways that influence growth regulation under stress.

To investigate potential synergistic functions between gh18-10 and nag-1, we attempted to generate a *nag-1* deletion in the Δ*gh18-10* background. Gene replacement was performed using a construct containing 1 kb homologous flanking regions surrounding an *erg27* resistance cassette and introduced into Δ*gh18-10* conidia by electroporation [29, 30]. Although fenhexamid-resistant fungal colonies were obtained, no viable double mutant could be identified, suggesting that the combined deletion of *gh18-10* and *nag-1* is lethal in *N. crassa*. Additionally, to determine whether Nag-1 functions as an active de-N-glycosylating enzyme, we attempted heterologous expression in *Pichia pastoris* and *Escherichia coli*. In both systems, the protein was expressed but not secreted, and it was detected only in the insoluble cell fraction.

In *Trichoderma asperellum*, two secreted β-N-acetylglucosaminidases (EXC1Y and EXC2Y) coexist, and disruption of *exc2y* does not impair vegetative growth, supporting functional overlap and redundancy among GlcNAcase activities [31]. Moreover, these enzymes are strongly induced by chitin-derived signals such as glucosamine. The dominant hexosaminidase can accumulate intracellularly in an active form before being secreted, indicating regulated mobilization of GlcNAcase capacity in response to metabolic cues [31]. Consistent with this, Nag1 from *Trichoderma reesei* is a functional GH20 enzyme that efficiently converts chitin oligosaccharides into GlcNAc, demonstrating that inducible NAGase activity can directly modulate chitin-derived carbohydrate pools [32].

Overall, these findings support a model in which upregulation of *nag-1* establishes an alternative biochemical and regulatory route. This pathway may partially compensate for disruptions associated with GH18, particularly the gh18-10 ENGase. Such compensation is likely most relevant under conditions of intensified glycan metabolism or elevated chitin- and amino-sugar–related signaling, including GlcNAc and glucosamine (GlcN). The coordinated regulation of these processes suggests that NAGase activity contributes to maintaining cellular homeostasis during stress, enabling adaptive responses to fluctuations in glycan turnover. By buffering perturbations in gh18-10–mediated functions, NAGase may help preserve cell wall remodeling dynamics, signaling balance, and structural integrity when canonical glycan-processing pathways are compromised. In summary, these results support the hypothesis that *N. crassa* might induce an alternative repertoire of enzymes with related catalytic architecture to mitigate ER stress when the primary de-N-glycosylating machinery is compromised.

## Materials and methods

### Strains and media

*Neurospora crassa* mutant and wild type (WT) strains used in the current study were provided by the Fungal Genetic stock center (Department of Plant Pathology, Kansas State University), and listed in Table S1. Strains were grown in Vogel’s N media [33], and incubated at 30°C in darkness.

### Protein expression and enzymatic activity assays

For heterologous expression in *S. cerevisiae png1Δ* cells, the *N. crassa* genes *gh18-10*, *gh18-11*, and *pngA* were amplified from cDNA and cloned into the pENTR™/D-TOPO vector following the manufacturer’s instructions (Invitrogen). Entry clones were recombined into the Gateway destination vector pYES-DEST52 and verified by DNA sequencing. Yeast transformation was performed as described previously [34]. Transformants were selected on SC-Ura agar plates containing 2% glucose and incubated at 30 °C.

Yeast cells, heterologously expressed the gh18-10, gh18-11 and pngA proteins, were grown overnight at 30 °C with shaking in SC-Ura-His medium supplemented with 2% raffinose. Expression driven by the GAL1 promoter was induced by transferring cells to SC-Ura-His medium containing 2% galactose and incubating for 6 h at 30 °C. Protein extraction was performed as described previously [35]. Briefly, approximately 1 × 10⁸ yeast cells were harvested, resuspended in 0.1 M NaOH, and incubated for 10 min at room temperature. Cells were collected by centrifugation at 3,000 × g for 1 min, and the pellets were resuspended in 100 µL SDS sample buffer (62.5 mM Tris– HCl, pH 6.8, 5% β-mercaptoethanol, 2% SDS, 5% sucrose, and 0.04% bromophenol blue). Samples were heated at 95 °C for 5 min, and cell debris was removed by centrifugation at 12,000 × g. Proteins were resolved by 15% SDS–polyacrylamide gel electrophoresis and confirmed by Western blot, using the mouse anti-V5 monoclonal antibody (Invitrogen), followed by incubation of the goat horseradish peroxidase-linked anti-mouse IgG secondary antibody (Rockland), and visualized ImmobilonTM Western blot chemiluminescent horseradish peroxidase substrate according to manufacturer’s instructions (Millipore). For protein deglycosylation, 10 µl of each sample was incubated with 2.5 mU of Endo H (Roche) in a total reaction volume of 20 µl at 37 °C overnight. Membranes were also probed with an anti-Dpm1 antibody (5C5, Invitrogen; 1:10,000 dilution) as a loading control.

In vitro deglycosylation activity of gh18-10, gh18-11 and pngA was assessed using S-alkylated RNase B [36]. Yeast cells expressing the above proteins were harvested, washed, and lysed in buffer containing 10 mM Tris–HCl, 1 mM EDTA, 250 mM sucrose, and 1 mM DTT, supplemented with 1× complete™ protease inhibitor cocktail and 1 mM Pefabloc (Roche). Lysates were prepared by glass-bead disruption followed by ultracentrifugation at 100,000 g for 1 h. Yeast extracts were incubated with S-alkylated RNase B overnight at 30 °C. RNase B treated with PNGase F (200 mU) served as a positive control, and S-alkylated RNase B incubated with lysis buffer as a negative control. gh18-10, gh18-11 and pngA -containing extracts were also treated with PNGase F to confirm ENGase-dependent band shifts. Deglycosylation was analyzed by immunoblotting using anti-RNase B antibody and HRP-conjugated secondary antibody. S-alkylated RNase B was prepared as described previously [36], with reduction performed in 10 mM DTT, 8 M urea, and 0.1 M Tris–HCl (pH 8.0).

### RTL spotting assay

The RTL assay was performed as described previously [22]. Briefly, *S. cerevisiae png1Δ* strains expressing RTL and *N. crassa gh18-10*, *gh18-11* and *pngA* were grown overnight at 30 °C with shaking in liquid SC-Ura-His medium supplemented with 2% raffinose. Four-fold serial dilutions of each culture were spotted onto SC-Ura-His-Leu and SC-Ura-His plates containing 2% galactose and incubated at 30 °C for 60h. A complemented *png1Δ* strain expressing *S. cerevisiae Png1* (*ScPng1*) and a strain carrying the empty *pRS316* vector served as positive and negative controls, respectively.

### Phenotypic stress analysis and crossings

For phenotypic stress analysis, *Neurospora crassa* strains (Table S1) were grown in Vogel’s N medium for 24 h at 30 °C. Fresh mycelial plugs (8 mm diameter) were excised using a cork borer and transferred to Vogel’s N medium supplemented with H₂O₂ (10 mM), SDS (0.008%), CoCl₂ (4 mM), NaCl (500 mM), caffeine (0.05%), DTT (2 mM), or tunicamycin (TNC; 0.005 mg/ml). Mycelial growth (diameter or perimeter) was measured after 24 h of incubation at 30 °C using ImageJ v1.54p from three biological replicates. Statistical analysis was performed using one-way ANOVA using the Tukey test (p < 0.05) (Minitab, LLC, 2021).

Crosses were performed using a modified previously published protocol [37]. Briefly, crossing plates were prepared by placing four 3.5-cm gel blot paper squares in 5-cm Petri dishes and adding 1 mL of synthetic crossing medium [38] (0.1% sucrose, 0.5% sorbose, 0.1% yeast extract, and 0.1% casamino acids). Each strain was first established as protoperithicial strain by evenly distributing 200 µL of a conidial suspension (1 × 10⁶ conidia in deionized water) onto the paper surface, followed by incubation in darkness at 25 °C for 48 h. Fertilization was then performed by adding 200 µL of a second 1 × 10⁶ conidia suspension onto the developing female mycelia, with these conidia serving as male gametes. Plates were incubated at 25 °C in darkness for 20 days, with brief light exposure during routine inspections. Ascospores were collected from plate lids using 1 mL of sterile deionized water, heat-treated at 65 °C for 1 h, plated onto selective media, and subsequently counted.

### RNA extraction and sequencing

Based on the phenotypic characterization, three stress conditions (ER, hypoxic and oxidative stress) were selected for subsequent transcriptomic analysis. *Neurospora crassa* strains were grown in liquid Vogel’s N broth for 12 h at 30 °C under continuous shaking (150 rpm). Subsequently, the mycelia were transferred to Vogel’s N broth supplemented with hydrogen peroxide (H₂O₂, 10 mM), cobalt chloride (CoCl₂, 4 mM), or tunicamycin (TNC, 0.005 mg/mL) and cultivated for an additional 24 h under the same conditions.

Mycelia were harvested, gently blotted with sterile wipes, and lyophilized to remove residual moisture. Total RNA was extracted using the Spectrum™ Plant Total RNA Kit (Sigma-Aldrich) according to the manufacturer’s instructions, and 1 μg of total RNA was treated with DNase I (Thermo Fisher Scientific). Strand-specific RNA-seq libraries were prepared using the TruSeq Stranded mRNA Library Preparation Kit with poly(A) selection (Illumina). Libraries were sequenced on an Illumina NovaSeq 6000 platform by Novogene. Three biological replicates were analyzed per treatment.

### RNA-seq data analysis

Raw sequencing reads were subjected to quality control using FastQC [39]. Read preprocessing was performed with PRINSEQ to filter sequences based on read length, GC content, quality scores, and duplicate reads. The *Neurospora crassa* OR74A genome (NCBI BioProject: PRJNA13841) was used as the reference genome and gene model annotation. Filtered paired-end reads were imported into Geneious Prime (v. 2025.0.4).

Read mapping to the reference genome was carried out using the Geneious RNA Mapper with the following parameters: minimum mapping quality of 30 bp, maximum gap per read of 5%, minimum overlap identity of 80%, and a minimum intron support of two reads. Gene expression levels were quantified in Geneious Prime by counting reads mapped to each gene. RPKM values were subsequently calculated based on the number of reads mapped to each gene and the length of the mappable gene region [40].

Differential gene expression analysis was performed using the DESeq package. Gene expression levels were compared between untreated control samples (no stress) and subsequent stress treatments. Differentially expressed genes were identified using a significance threshold of P < 0.05 and a log₂ fold change greater than 2 or less than −2. Functional annotation and Gene Ontology (GO) enrichment analyses were conducted using Blast2GO [41]. Expression heatmaps and GO bubble plots were generated using OmicsBox and g:Profiler (BioBam Bioinformatics). Venn diagrams were constructed using Venny 2.1. PDB structures were generated using AlphaFold 3 [42]. Structural alignments were performed with TM-align using default parameters [43].

## Acknowledgements

This work was supported by the Carl Tryggers Foundation (grant no. CTS22:1920), the Helge Axelsson Johnson Foundation (grant no. F25-0341), the Royal Swedish Academy of Agriculture and Forestry (KSLA) (grant no. GFS2024-0151), the RIKEN International Joint Graduate School Program, Japan, and the Swedish University of Agricultural Sciences. We sincerely thank Prof. Tadashi Suzuki (Glycometabolic Biochemistry Laboratory, RIKEN) for hosting us in his laboratory and for his valuable contributions, guidance and mentorship to the expression and characterization of the *N. crassa* enzymes.

## Author contributions

GT and MK conceived the project, AS, GT and TJH designed and executed the experiments, AS and GT analyzed the data, AS and GT wrote the first draft of the manuscript. All authors contributed to the editing of the manuscript and have been read and agreed to the final version of the manuscript.

## Data availability statement

The data support this study are available in the Material and Methods and in the Supplementary Information of this article. The transcriptomic data of this study were deposited in the NCBI database (https://www.ncbi.nlm.nih.gov/) under BioProject accession number PRJNA1427682.

