## Supplementary files for "Functional and transcriptomic analyses in *Neurospora crassa* reveal the crucial role of N-glycoprotein deglycosylation process in fungal homeostasis"

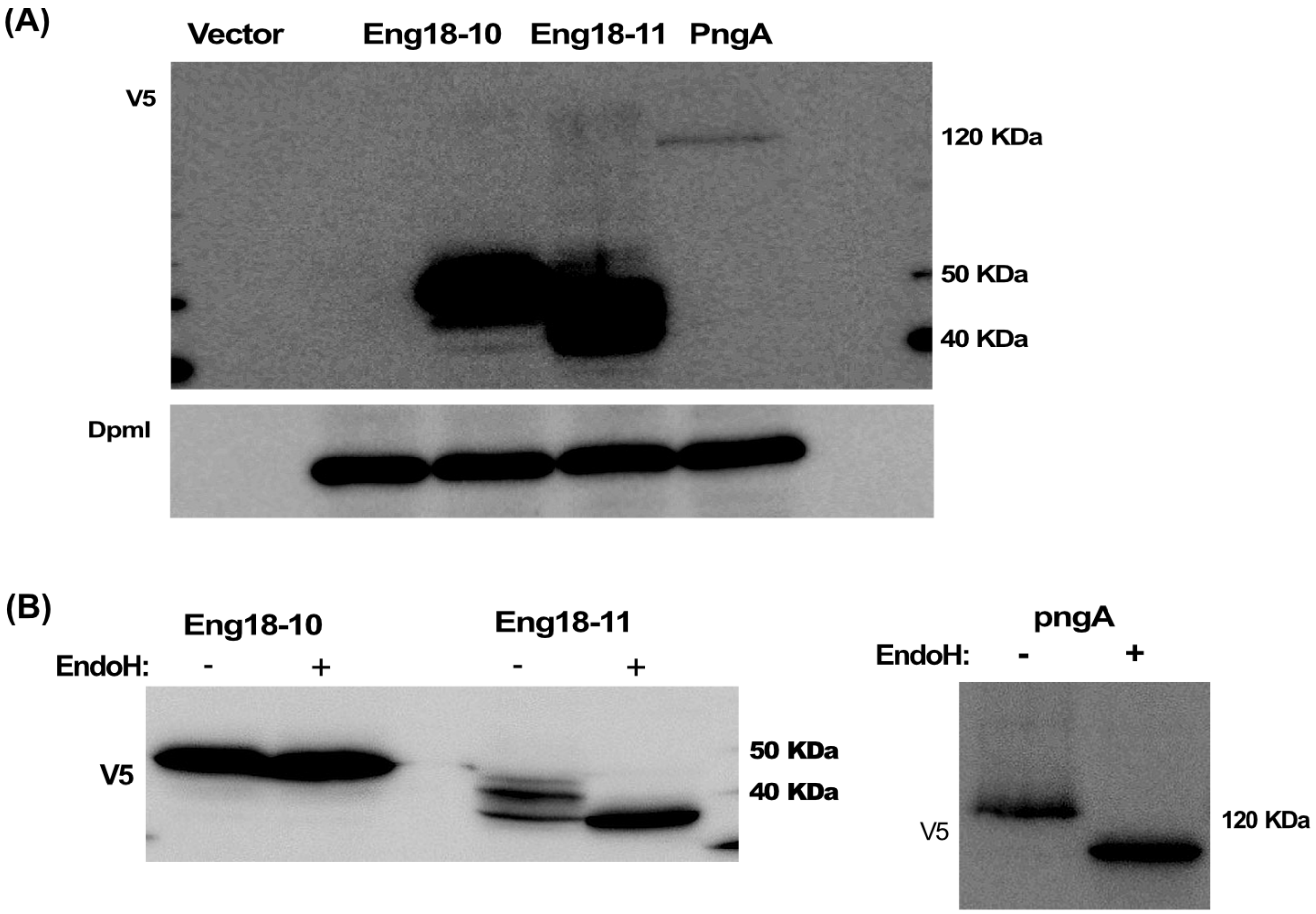


**Fig. S1.** Heterologous expression of *Neurospora crassa* ENGases and PNGase in *Saccharomyces cerevisiae*. (A) *Neurospora crassa* gh18-10, gh18-11 and pngA were heterologously expressed in png1Δ cells. Treatment of membrane with the anti-Dpm1 antibody used as a loading control.  Yeat cells expressing the pYES-DEST52 empty vector was used a negative control (B) Treatment of gh18-10, gh18-11 and pngA with (+) or without (-) Endo-H. Proteins were resolved by 15% SDS–polyacrylamide gel electrophoresis and confirmed by Western blot, using the mouse anti-V5 monoclonal antibody, followed by incubation of the goat horseradish peroxidase-linked anti-mouse IgG secondary antibody.

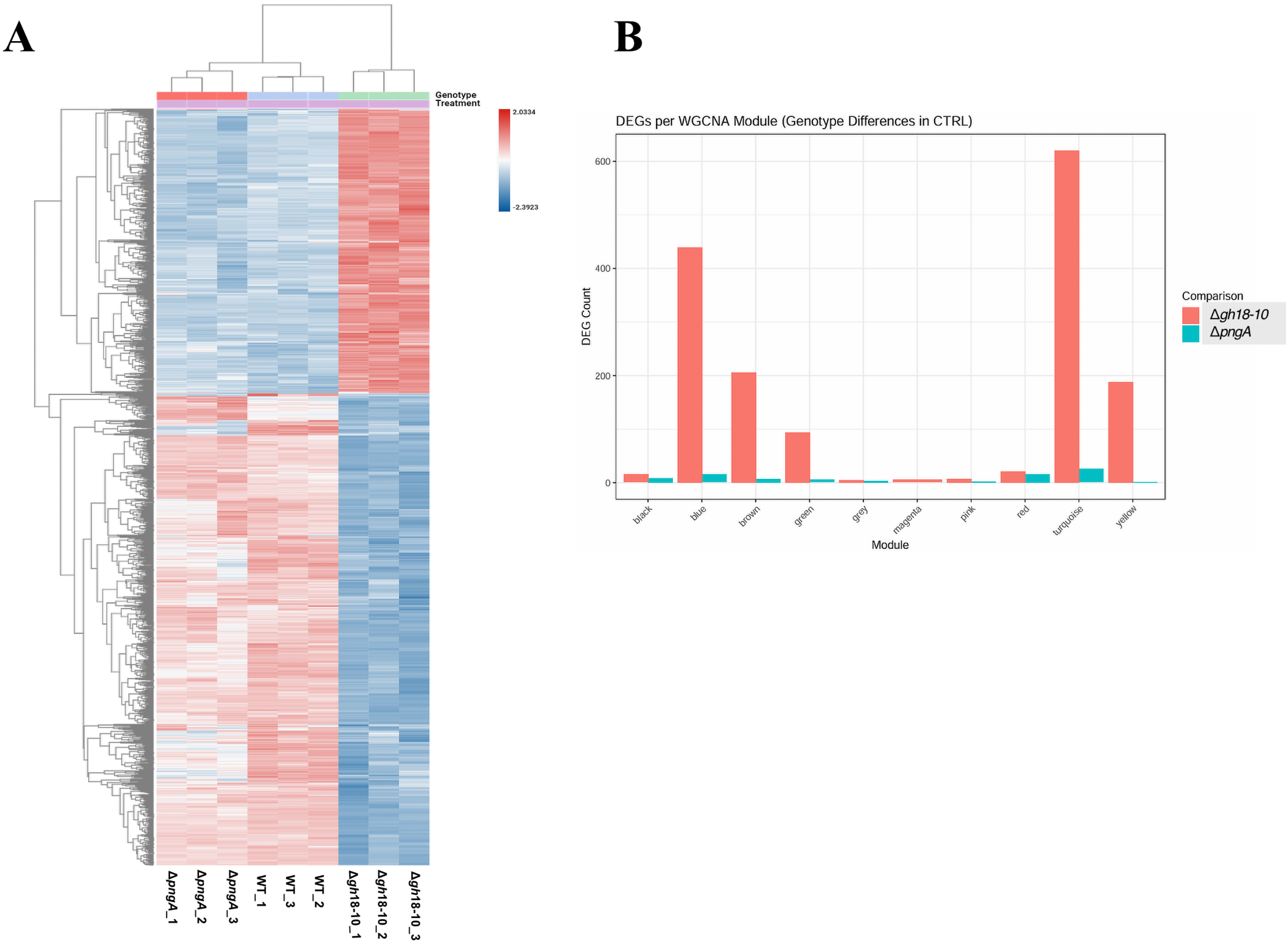


**Fig. S2.** (A) Heatmap of differentially expressed genes (DEGs) (p.value=0,05 and -2<log2fold>2) comparing WT with the Δ*gh18-10* and Δ*pngA* mutants under control conditions. Expression values are shown as row-scaled, normalized transcript levels, and samples are clustered based on similarity. (B) Weighted gene co-expression network analysis (WGCNA) of control-responsive genes depicting the number of DEGs per co-expression module for each genotype (Δ*gh18-10* and Δ*pngA*).

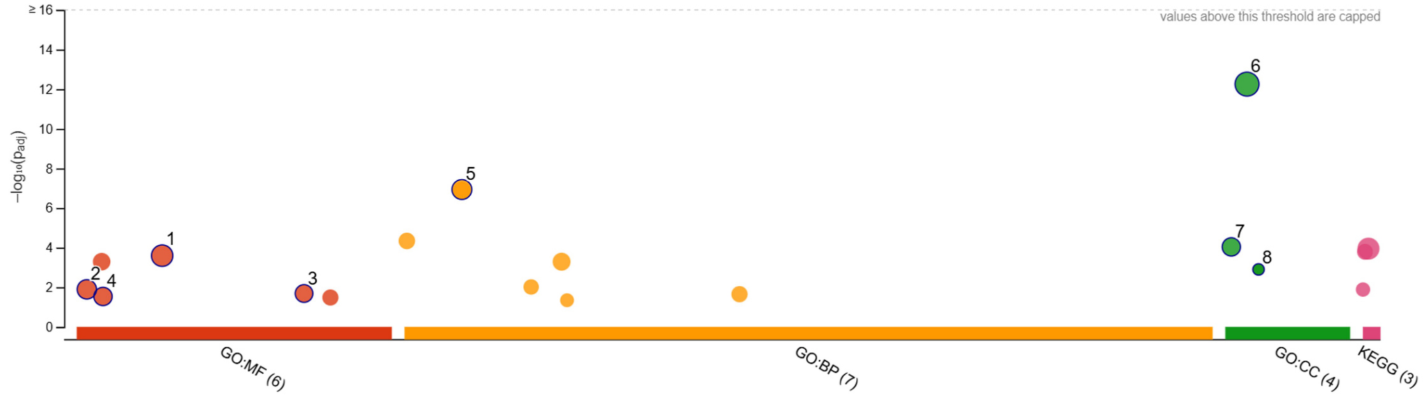


| ID | Source | Term ID | Term Name | $p_{adj}$ (query_1) |
|---|---|---|---|---|
| 1 | GO:MF | GO:0016491 | oxidoreductase activity | $2.546\times10^{-4}$ |
| 2 | GO:MF | GO:0003700 | DNA-binding transcription factor activity | $1.278\times10^{-2}$ |
| 3 | GO:MF | GO:0050660 | flavin adenine dinucleotide binding | $2.070\times10^{-2}$ |
| 4 | GO:MF | GO:0004553 | hydrolase activity, hydrolyzing O-glycosyl compou... | $2.920\times10^{-2}$ |
| 5 | GO:BP | GO:0005975 | carbohydrate metabolic process | $1.163\times10^{-7}$ |
| 6 | GO:CC | GO:0016020 | membrane | $5.539\times10^{-13}$ |
| 7 | GO:CC | GO:0005576 | extracellular region | $9.185\times10^{-5}$ |
| 8 | GO:CC | GO:0031380 | nuclear RNA-directed RNA polymerase complex | $1.260\times10^{-3}$ |

| Gene Names | Protein names | Gene Ontology IDs | Biological Terms |
|---|---|---|---|
| **NCU02925** | FAD-dependent monooxygenase srdH (Fruiting body maturation protein 1) (Sordarial biosynthesis cluster protein srdH) | GO:0004497; GO:0044550; GO:0071949 | fruiting body |
| **NCU09915** | Female sexual development-1 protein, variant 1 | GO:0003677; GO:0003700 | development |
| **NCU01510** | Meiotically up-regulated 190 protein | GO:0005789; GO:0006869; GO:0008289; GO:0061817 | meiotic |
| **NCU08064** | Meiotically up-regulated 190 protein | GO:0005789; GO:0006869; GO:0008289; GO:0061817 | meiotic |

**Fig. S3.** g:Profiler enrichment analysis of genes downregulated in Δ*gh18-10* as compared to WT under control conditions. The bubble plot summarizes significantly enriched terms across GO categories (molecular function, biological process, cellular component) and KEGG; the y-axis indicates enrichment significance (−log10 adjusted p-value). Bubble size reflects the number of genes associated with each term. The table lists the top enriched terms and corresponding adjusted p-values, and the lower panel highlights representative downregulated genes annotated with sexual development–related terms.

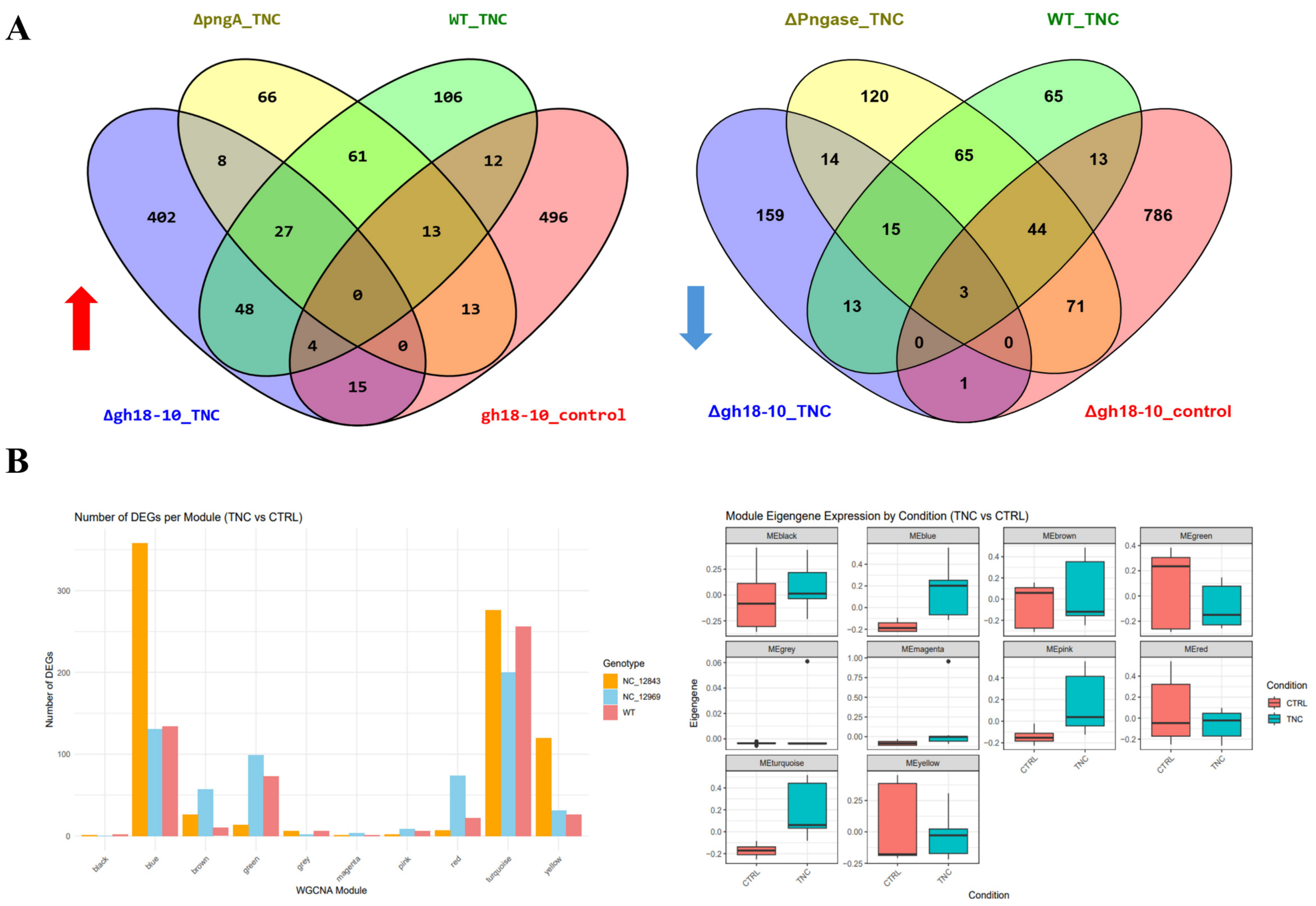


**Fig. S4.** (A) Venn diagrams showing the overlap of differentially expressed genes (DEGs) under tunicamycin (TNC) stress treatment relative to Δ*gh18-10* control conditions in the indicated genotypes. The left panel (red arrow) represents upregulated DEGs, and the right panel (blue arrow) represents downregulated DEGs. Numbers indicate the count of shared or unique DEGs in each intersection. (B) Weighted gene co-expression network analysis (WGCNA) of TNC-responsive genes. The left bar plot shows the number of DEGs per co-expression module (TNC vs CTRL) for each genotype. The right panels display boxplots of module eigengene expression across conditions (CTRL vs TNC).

**A**

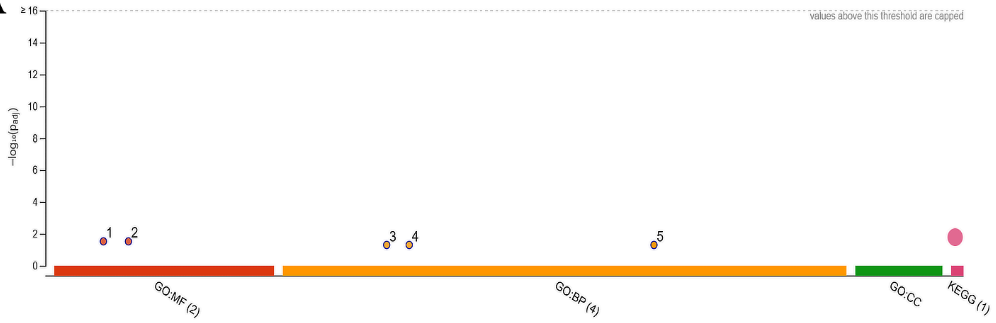


| ID | Source | Term ID | Term Name | $p_{adj}$ (query_1) |
|---|---|---|---|---|
| 1 | GO:MF | GO:0015106 | bicarbonate transmembrane transporter activity | $2.988\times10^{-2}$ |
| 2 | GO:MF | GO:0019531 | oxalate transmembrane transporter activity | $2.988\times10^{-2}$ |
| 3 | GO:BP | GO:0015701 | bicarbonate transport | $4.993\times10^{-2}$ |
| 4 | GO:BP | GO:0019532 | oxalate transport | $4.993\times10^{-2}$ |
| 5 | GO:BP | GO:0072383 | plus-end-directed vesicle transport along microtu... | $4.993\times10^{-2}$ |

**B**

| UniProt ID | Gene | Protein | Biological Process |
|---|---|---|---|
| **Q7S6E8** | gh81-1 NCU07076 | glucan endo-1,3-beta-D-glucosidase (EC 3.2.1.39) | cell wall organization [GO:0071555]; polysaccharide catabolic process [GO:0000272] |
| **Q7RV09** | ars-1 NCU06041 | Arylsulfatase (AS) (EC 3.1.6.1) (Aryl-sulfate sulphohydrolase) | phenol-containing compound metabolic process [GO:0018958] |
| **V5IPM4** | NCU10852 | Beta-hexosaminidase (EC 3.2.1.52) | carbohydrate metabolic process [GO:0005975]; glycosaminoglycan metabolic process [GO:0030203] |
| **U9W500** | NCU00244 | Glycosyl transferase | |

**Fig. S5.** (A) g:Profiler functional enrichment summary for the Δ*gh18-10* tunicamycin (TNC)–specific upregulated gene set. Dots represent significantly enriched terms; the y-axis indicates enrichment significance (adjusted p-value scale), and the table lists the top enriched categories. (B) Selected Δ*gh18-10* TNC–specific upregulated genes with annotations relevant to glycosylation processes.
